# A preliminary investigation on the dependence of arthropod diversity on vegetation diversity across four contrasting ecosystems in Hanthana mountain range of Sri Lanka

**DOI:** 10.1101/637280

**Authors:** W.A.M.T. Weerathunga, A.M.G.K. Athapaththu, L.D. Amarasinghe

## Abstract

Arthropods contribute significantly to biodiversity and vegetation provides a habitat and resources for them to survive, exist and propagate. We report a preliminary investigation on the extent to which arthropod diversity is dependent upon vegetation diversity across different ecosystems in a humid tropical climate. We determined the diversity of arthropods in four ecosystems closely-located ecosystems with different vegetation. Vegetation surrounding an aquatic environment (AQ), a broad-leaved wet, evergreen forest ecosystem (BL), a *Pinus caribaea* monoculture plantation (PN) and a *Pinus* plantation artificially enriched with indigenous broad-leaved tree species (PNEN) located in the Hanthana mountain range in Central Sri Lanka were selected. In each environment, arthropods were sampled in three randomly-selected sites (5 m x 5 m) using four sampling methods. Collected arthropods were identified upto the highest possible taxa using standard identification keys. Simultaneously, vegetation diversity was determined via a plant census. Arthropod and vegetation diversities were computed separately for each site using Shannon-Wiener Index (H).

Within the 300 m^2^ area of observation plots, arthropod individuals belonging to 68 species and 43 families were found. AQ had the greatest arthropod diversity (H=2.642), dominated by *Olios* spp. followed by BL (H=2.444), dominated by three arthropods, namely, a tettigonid species, *Oxytate* spp. and *Psechrus* spp. PN had the next highest arthropod diversity (H=1.411), dominated by *Dicaldispa* spp. The lowest arthropod diversity was found at PNEN (H=1.3500), dominated by an ant species. Contrastingly, PNEN had the highest plant diversity (H=2.614) and PN the lowest (H=0.879). AQ (H=1.810) and BL (H=1.871) had intermediate values.

In a regression involving data from AQ, BL and PN, arthropod diversity was linearly dependent on plant diversity (R^2^=0.423) whereas it was not so when PNEN was also included (R^2^=0.008). This finding supports the hypothesis that while higher plant diversity contributes to greater arthropod diversity in ecosystems where human intervention is minimal, artificial enrichment of plant diversity does not necessarily increase arthropod diversity in the short-to medium-term. Further investigations are needed to substantiate these preliminary findings and validate the above hypothesis.

## Introduction

Phylum Arthropoda has been recognized as the largest phylum in the animal kingdom as it alone accounts for about 80 % of the total number of described species (Zhang, 2011). Arthropods are a highly versatile group of organisms being the first to make a move into air, successfully colonizing in all three media-land, water and air. Like all other animals, arthropods are also known to have a strong interrelationship with their surrounding vegetation, with herbivorous arthropods and their associated trophic levels, playing a major role. Several studies have identified plant diversity as an important determinant of arthropod diversity (May, 1978; Hunter & Price, 1992; Haddad *et al.*, 2009). Several models (MacArthur, 1972; Whittaker, 1975; Tilman, 1980; Rosenzweig, 1995) and experimental studies (Alteri & Letourneau 1982; Lawton, 1983; Siemann *et al*., 1998; Bassett et al., 2012; Castangneyrol & Jactel, 2012) bear evidence for increase of arthropod diversity with increasing plant diversity. There are two major hypotheses to explain this phenomenon, namely Resource Specialization Hypothesis (RSH) and More Individual Hypothesis (MIH). According to RSH, approximately 90% of herbivorous arthropods show host specific specialization (Bernays & Graham, 1988). Therefore as the plant species richness increases the number of associated herbivore species should also increase accordingly (Hutchinson, 1959; MacArthur, 1972; Price *et al*., 1980). In contrast, MIH states that if above-ground net primary productivity (ANPP) increases as plant species richness increases then more herbivore individuals will be supported owing to the increased availability of resources (Srivastava & Lawton, 1998; Hooper, 2005). Therefore, an increased number of herbivore species by either of these hypotheses shall support more predator species (Siemann *et al*., 1998; Srivastava & Lawton, 1998; Haddad *et al*., 2001; Crutsinger *et al*., 2006; Johnson *et al*., 2006).

Influence of vegetation diversity may be different for diversity of arthropods in different species groups and trophic levels. For example, Korichova et al. (2000) showed that vegetation diversity influenced the abundance of only the most sessile and specialized arthropod taxa. On the other hand, the abundance of leafhoppers decreased linearly with increasing plant species richness. Stronger positive correlations between arthropod- and vegetation diversity have been shown for primary-than secondary consumers (Castangneyrol & Jactel, 2012). On the other hand, Haddad *et al.* (2009) showed that species richness of both herbivore and predator arthropods were strongly positively correlated to plant species richness. Notably, Haddad *et al.* (2009) found that arthropod species richness shifts from a predator-dominated trophic structure to a herbiover-dominated structure with decreased plant species richness. Interestingly, diversity of herbivore arthropods has been shown to be more strongly related to diversities of predators and parasites than to plant diversity (Siemann *et al.*, 1998). Also of interest is the finding of Bangert *et al.* (2005) that arthropod diversity increases with increasing genetic diversity within a given plant species. Furthermore, Wimp *et al.* (2005) showed that arthropod community composition of cottonwood responded to the genetic variation among its host tree community. This suggests that increasing genetic diversity in the vegetation community is a pathway of conserving the diversity of its arthropod community.

Architectural or structural diversity of plants, probably correlating with their functional and species diversity could determine arthropod diversity (Lawton, 1983; Brose, 2003; Woodcock *et al.*, 2010). Meanwhile, changing plant diversity can play a major role in interactions between herbivorous arthropods and their predators and parasites (Pimentel, 1961; Strong *et al*., 1984; Andow & Prokym, 1990; Coll & Bottrel, 1996; Haddad *et al.*, 2009). Vegetation diversity can vary in three major ways namely the phylogeny, spatial array, and the temporal overlap of plants in the mixture (Andow, 1991). Agricultural and landscaping activities actively influence these variations attributing to the growing concern of biodiversity decline in agricultural landscapes (Matson *et al*., 1997; Krebs *et al*., 1999; Yaacobi *et al.*, 2007; Isaacs *et al.*, 2009).

Hanthana mountain range (7° 15’ N, and 80° 37’ E), the study area of the present research, has been declared as an environmental protection area (Gazette Notification No.1641/28, Environmental Protection Areas, 2013). It is located within the humid tropical climatic zone in the Central Highlands of Sri Lanka and traverses an altitudinal range from *ca.* 500 to 800 m above mean sea level, displaying substantial spatial variation in plant and ecosystem diversity. The different ecosystems present within the Hanthana mountain range include tropical broadleaved wet evergreen forests, interspersed with grasslands and monoculture *Pinus carribaea* plantations. Historical evidence suggests that broadleaved wet evergreen forests were the dominant vegetation type in this area. During the 18^th^ and 19^th^ centuries, part of this forest had been cleared to establish plantations of tea (*Camellia sinensis*) and rubber (*Hevea brasiliensis*). Improper management of these cropping systems caused substantial soil erosion and consequent land degradation, which made the tea and rubber plantations economically non-viable by mid-20^th^ century. This led to these lands being abandoned in a highly-degraded state with only a perennial grassland cover dominated by *Panicum maximum* and *Cymbopogon nadus*, thus posing a serious threat to the ecosystem stability of the entire area.

In the 1960s and 70s, *Pinus* was planted in these grasslands as a means of establishing a perennial tree cover to arrest soil erosion and allow gradual development of a natural forest ecosystem via the process of succession. Principal reasons for selection of *Pinus* for reforestation of the grasslands were its ability to establish in the degraded infertile soil and tolerate periodic fires that occur during dry periods. However, the expected natural succession of *Pinus* plantations did not take place as very few plant species could penetrate and colonize the thick mat of pine needles on the ground, which is highly-resistant to decomposition. As a means of restoring the soil and vegetation and increasing biodiversity, an initiative was taken during the early 1990s to plant selected indigenous tree species by partial removal of some *Pinus* plants and using the shade of the remaining *Pinus* plants as a ‘nurse crop’ to aid establishment of the newly-introduced plant species (Ashton *et al.*, 1997). This has been successful in the limited area that it was undertaken. The indigenous tree species have established among the remaining *Pinus* plants, thus giving rise to a mixed forest plantation. The process of natural succession appears to be taking place slowly and vegetation diversity has increased. Numerous work elsewhere in the world, where vegetation diversity had been increased or decreased artificially, report subsequent increases or decreases in arthropod diversity <u>(Knops et al. 1999; Wyss 1996)</u>. However, there has been no previous work to determine whether the same has happened in *Pinus* plantations ‘enriched’ with indigenous tree species (termed ‘enriched *Pinus*, PNEN) in the Hantana mountain range. In fact, except for studies targeting a particular group of arthropods (Chathuranga & Ranawana, 2017), a general survey on arthropod diversity has not been carried out in this area in the recent past.

Therefore, in the present work, which was intended to be a preliminary short-term investigation, our objectives were to find answers to the following questions: (a) Do the different ecosystems that are present within the Hanthana mountain range show significant variation in their arthropod diversity?; (b) If so, is there evidence to support the generally established positive relationship between arthropod diversity and vegetation diversity? (c) Has the artificial increase of vegetation diversity in the monoculture *Pinus* plantations via enrichment planting of indigenous tree species resulted in an increase in arthropod diversity after two decades?

Furthermore, the Hanthana mountain range has been an area of severe human intrusion during the past four decades. As it is located close to a major city (Kandy, Central Sri Lanka) and has several villages and tea plantations in its borders, encroachment and subsequent deforestation is continuing for expansion of both residential areas and tea plantations. While the impact of such activities on vegetation diversity has been obvious, there are no previous studies on the impacts on the diversity of arthropods, a key faunal group, with numerous essential roles in sustaining biodiversity and ecosystem stability. We believe that the present work will provide valuable information, which can be part of plans and programmes to conserve the biodiversity and ecosystem stability in the Hanthana mountain range.

## Methodology

### Study Area

Hanthana mountain range (7° 15’ N, and 80° 37’ E) is located in the mid-country wet zone of Sri Lanka, and is divided into two major regions, namely Upper Hanthana mountain area (> 600 m) and Lower Hanthana mountain area (< 600 m). It has been identified that the Lower Hanthana area is subjected to heavy human encroachment and Upper Hanthana area is comparatively pristine (Chathuranga & Ranawana, 2017). This study was carried out during the months of August-September, 2016 in a dry season following the South-Western monsoons.

### Experimental design

Four contrasting ecosystems located close to each other in Hanthana mountain range, were selected as different treatments of the study. The four ecosystems were namely, vegetation surrounding an aquatic environment (AQ), a broad-leaved, wet evergreen ecosystem (BL), *Pinus caribaea* monoculture vegetation (PN) and a *Pinus* plantation artificially enriched with broad-leaved tree species (PNEN). In each of these ecosystems three replicate sites (5 m x 5 m) were chosen randomly and temporarily demarcated. AQ and BL treatments were located in the Lower Hanthana area while PN and PNEN treatments were located in Upper Hanthana area (Fig. 1).

**Figure. 1.**
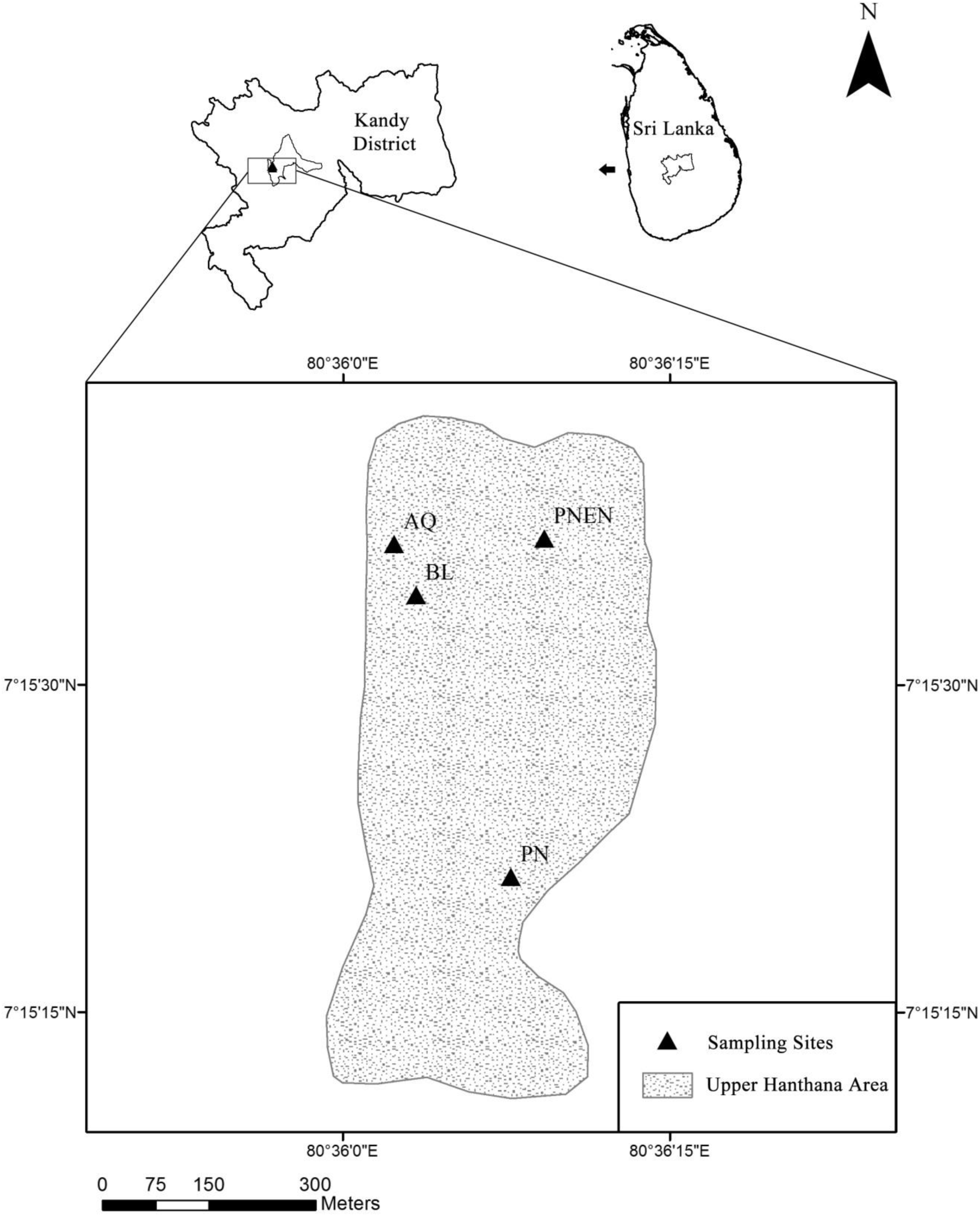
Map of study area showing sampling sites Aquatic Vegetation (AQ), Broad Leaved Vegeation (BL), *Pinus* monoculture (PN) and Enriched *Pinus* Vegetation (PNEN) AQ was a flatland bordered by a man-made water body on one side. BL ecosystem was an area with a slight inclination and a thick growth of broad-leaved wet evergreen tree species forming a dense canopy, in contrast to PN where *Pinus caribaea* was the dominant tree species with a *Panicum maximum* dominated grassland as undergrowth. PNEN was a land which had been a *Pinus caribaea* monoculture previously, but had been enriched with artificially recruited broad-leaved species about 20-25 years ago.

### Data collection on arthropods and plants

Arthropods in each 5 m x 5 m replicate site were sampled using four different sampling methods (i.e. pit fall traps, sticky traps, sweep net and beating tray) in a way that arthropods at all heights from ground level are covered. Around 40-50 sweeps were done by the sweep net in each replicate site, for sampling arthropods at moderate heights above ground. A circular cloth having a diameter of 120 cm was used as the beating tray, which was held under unreachable foliage in the sampling plot, for sampling arthropods at unreachable heights above ground. Self-designed pit-fall traps and sticky traps were set for a period of one week, for sampling arthropods at ground level. Pit-fall traps were prepared by cutting a plastic water bottle of diameter in half. A piece of cotton wool soaked in chloroform (0.5 mL) was placed at the bottom of each pit-fall trap and they were covered with a metal sheet (Fig. 2). Square plastic sheets (15 cm x 15 cm) spread with vaseline were used as sticky traps, for sampling arthropods at mid-level above ground. Three pit-fall traps and three sticky traps were used for each replicate site. The traps were left for a period of one week and the arthropods collected were preserved in 70% alcohol and were identified to the highest possible taxa using standard identification keys, based on their morphological characteristics. Simultaneously, a plant census was also done for each replicate site to identify the plant species and their abundance. Every plant that was present in the sampling plot was individually identified using their morphological characteristics, with the help of standard pictorial guides and herbarium specimens.

**Figure. 2:**
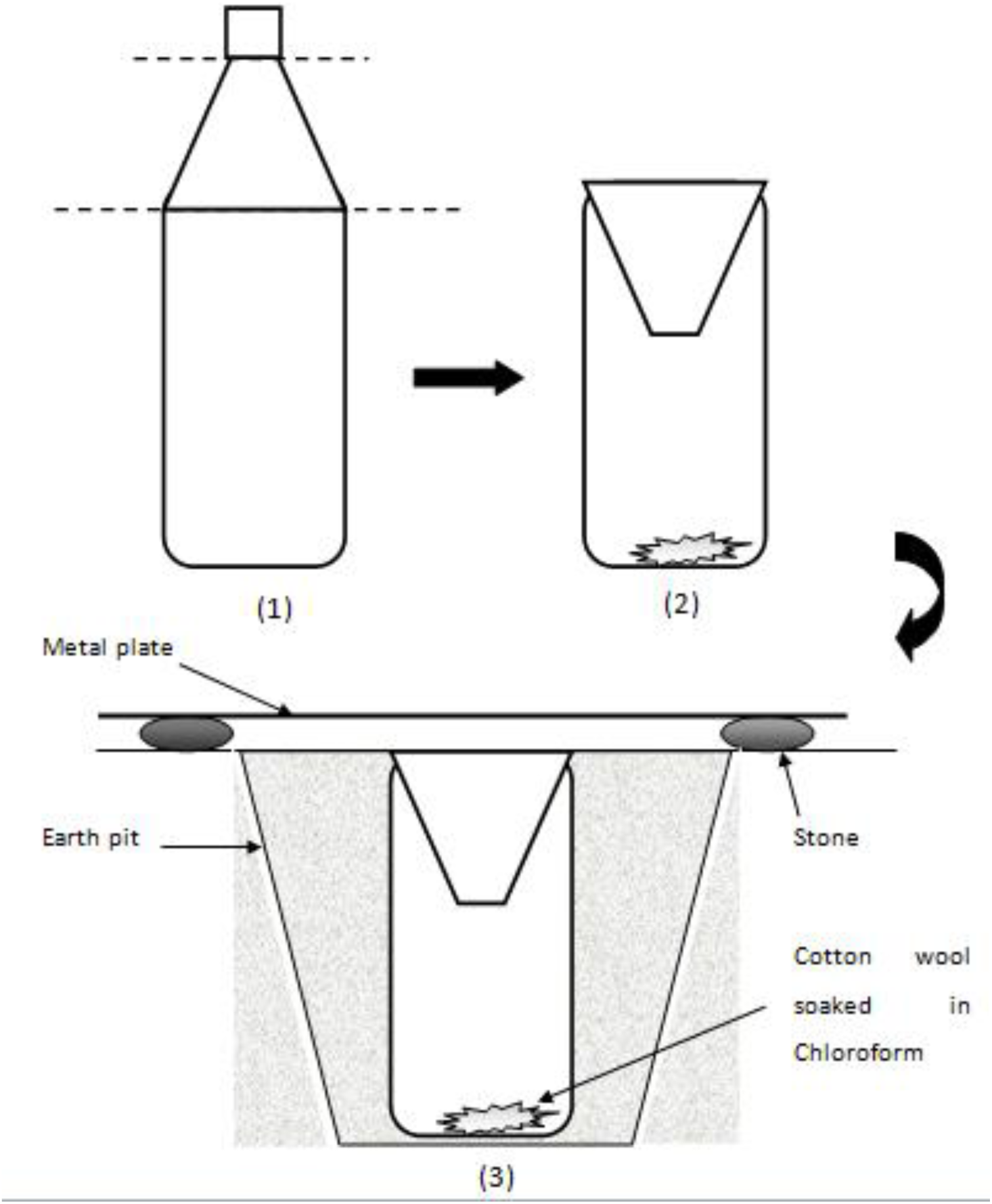
Schematic diagram of the way of preparing a pit-fall trap from a plastic water bottle and (2) and as established on the ground (3)

### Statistical Analysis

The diversity of arthropods and plants were calculated separately for each replicate site, using the Shannon-Wiener Index (H) along with species richness and evenness (Magurran, 2004). The significance of variation between the four ecosystems, based on arthropod diversity was determined by one-way analysis of variation (ANOVA) for the diversity indices obtained for each replicate site of each ecosystem. Mean separation was done by using the Least Significant Difference. The dependence of arthropod diversity on plant diversity was determined by conducting a simple linear regression analysis. The variation among the four ecosystems based on arthropod diversity and vegetation diversity was determined separately, using principle component analysis. All statistical analyses were carried out using Minitab 14.0 and Primer 5 software packages.

## Results

Arthropod individuals belonging to 68 species and 43 families were collected from the 12 sampling sites (5 m x 5 m) across the four ecosystems. In the same sampling area plant species belonging to 84 species and 42 families were enumerated.

Based on the Shannon-Wiener Index (H), the vegetation surrounding the aquatic environment (AQ) had the highest arthropod diversity (H=2.642) as well as the highest plant diversity. The most abundant arthropod found in this ecosystem was identified as *Olios* spp. (Araneae; Family Sparassidae) (Fig. 3). The second highest arthropod diversity (H=2.444) was found in the broad-leaved, wet, evergreen ecosystem (BL). It was dominated by three arthropods, namely, a tettigonid species, *Oxytate* spp. (Araneae: Family Thomisidae, a crab spider genus), and *Psechrus* spp. (Family Psechridaea jungle cribellate spider genus) (Fig. 4). *Pinus caribaea* monoculture vegetation (PN) had the third highest arthropod diversity (H=1.411) and it was dominated by *Dicaldispa* spp. (Coleoptera; Chrysomelidae) (Fig. 5). The lowest arthropod diversity (H=1.3500) was found in the Pinus plantation artificially enriched with broad-leaved species (PNEN), which was dominated by an ant species (Hymenoptera; Formicidae) (Fig. 6). In contrast, when considering plant diversity, PNEN had the highest diversity (H=2.614) and PN the lowest (H=0.879). AQ (H=1.810) and BL (H=1.871) had intermediate values.

**Figure. 3:**
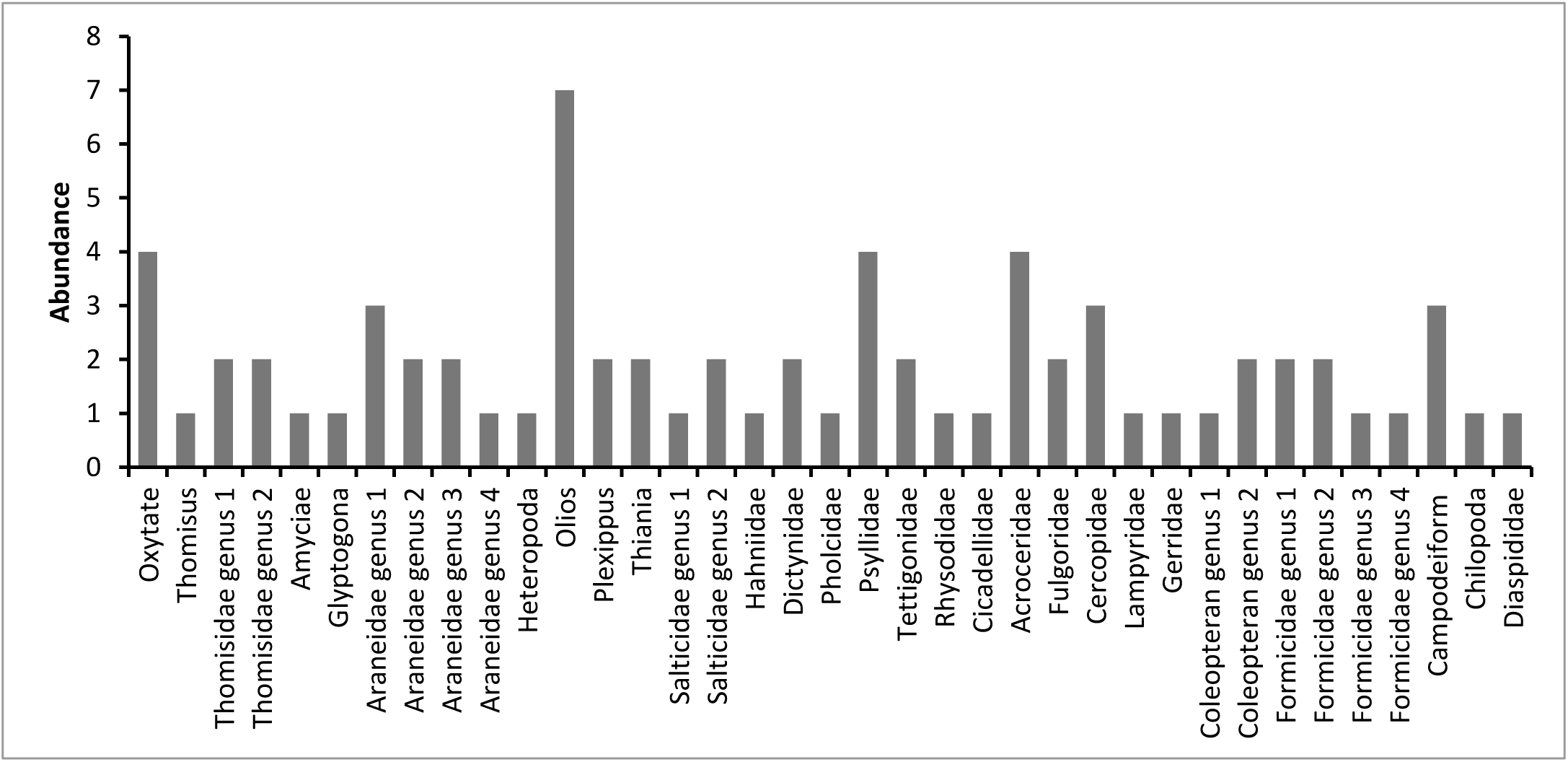
Arthropod diversity in Aquatic based vegetation (AQ).

**Figure. 4:**
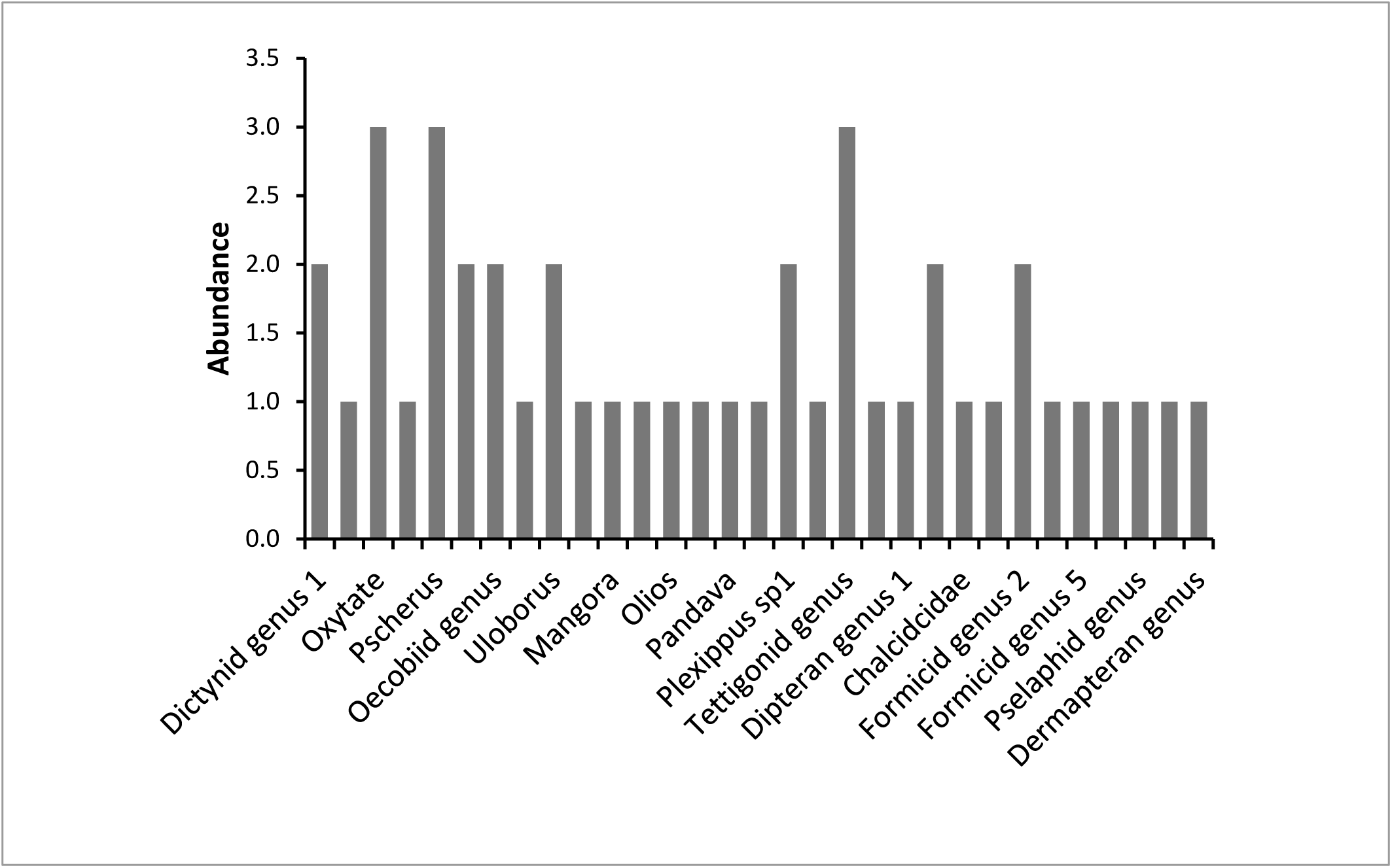
Arthropod diversity in Broad leaved vegetation (BL).

**Figure. 5:**
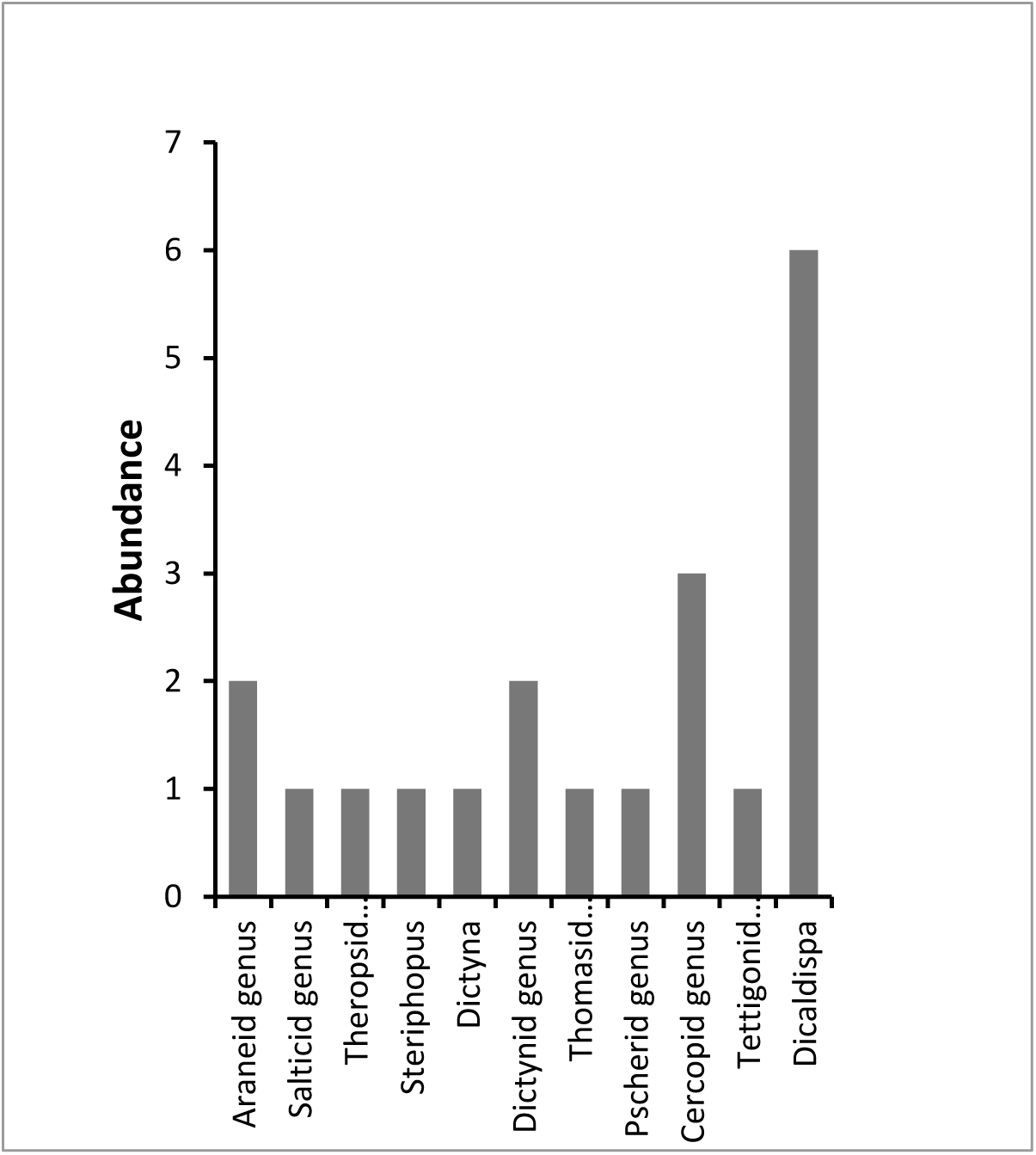
Arthropod diversity in *Pinus* monoculture (PN).

**Figure. 6:**
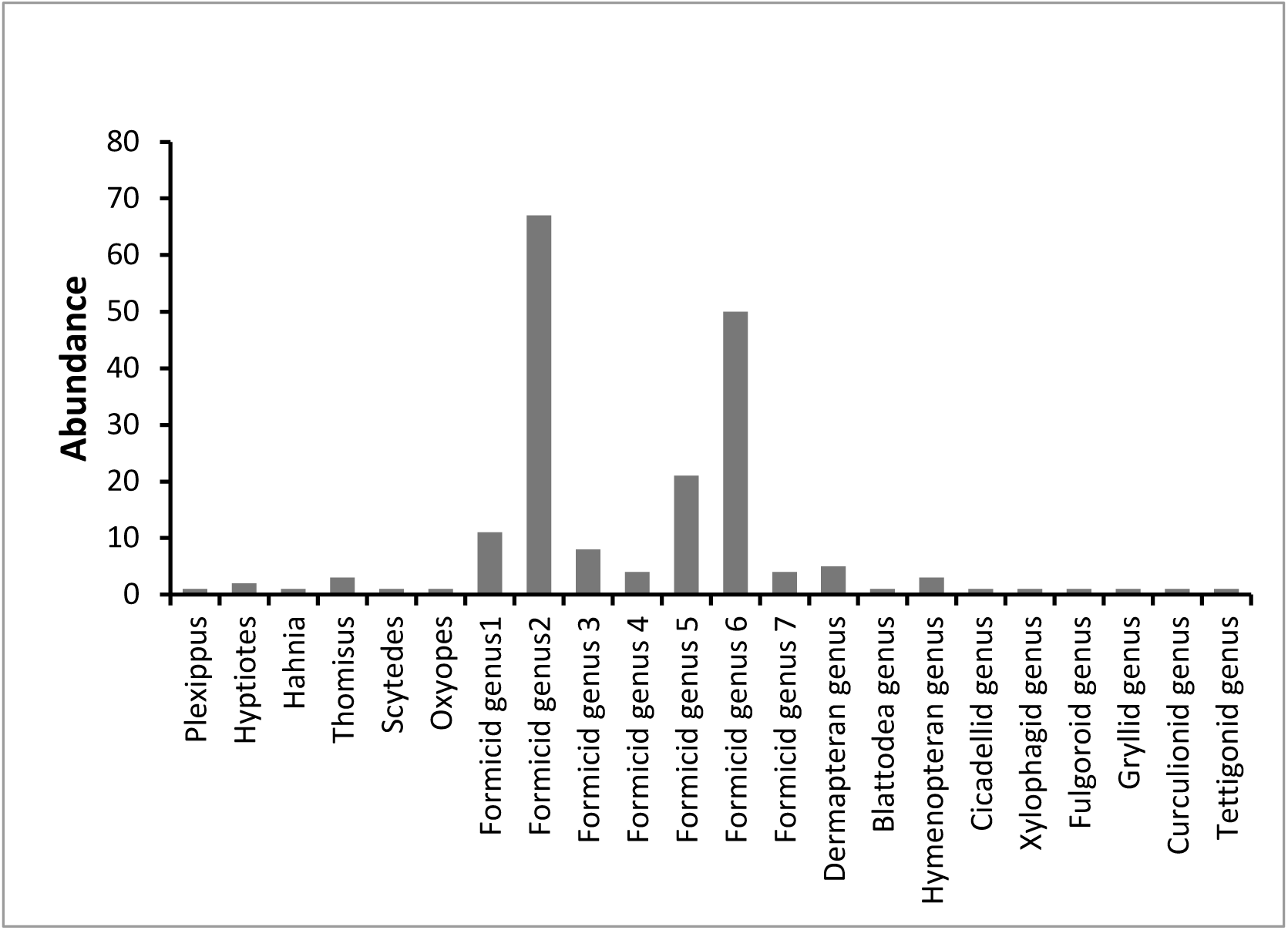
Arthropod diversity of Enriched *Pinus* vegetation (PNEN).

According to the ANOVA for Shannon-Wiener diversity indices of each ecosystem, arthropod diversity was significantly higher (*p*<0.1) in AQ and BL (Table 1) than in PN and PNEN. In contrast, the Shannon-Wiener Index for plant diversity was significantly (*p*<0.01) highest in the PNEN and was lowest in PN while AQ and BL had intermediate values. Arthropod species richness showed significant (*p*<0.001) variation among ecosystems with PN having a significantly lower value than the others, which did not differ significantly. In contrast, the ecosystems did not differ significantly in terms of plant species richness. Evenness of species distribution of both Arthropods and plants showed significant variation among ecosystems (Table 1).

**Table 1:** Variation of Arthropod- and vegetation diversity in the ecosystems selected for the study

| Ecosystem | Arthropods |  |  | Vegetation |  |  |
| --- | --- | --- | --- | --- | --- | --- |
|  | Shannon- | Species | Evenness | Shannon- | Species | Evenness |
|  | Wiener | Richness |  | Wiener | Richness |  |
|  | Index |  |  | Index |  |  |
| Aquatic | 2.641<sup>a</sup> | 16.33<sup>a</sup> | 0.972<sup>a</sup> | 1.810<sup>b</sup> | 22.33<sup>a</sup> | 0.601<sup>b</sup> |
| Broad-leaved | 2.444<sup>a</sup> | 12.67<sup>ab</sup> | 0.983<sup>a</sup> | 1.871<sup>b</sup> | 15.00<sup>a</sup> | 0.698<sup>b</sup> |
| *Pinus* | 1.411<sup>b</sup> | 5.00<sup>c</sup> | 0.974<sup>a</sup> | 0.879<sup>c</sup> | 21.67<sup>a</sup> | 0.286<sup>c</sup> |
| *Pinus* enriched | 1.350<sup>b</sup> | 9.67<sup>bc</sup> | 0.635<sup>b</sup> | 2.614<sup>a</sup> | 19.67<sup>a</sup> | 0.884<sup>a</sup> |
| CV (%) | 26.99 | 43.90 | 8.47 | 22.38 | 32.68 | 18.48 |
| $p>F$ | 0.057 | 0.0005 | 0.0032 | 0.011 | ns | 0.0038 |

Along each column, means with the same letter are not significantly different at $p = 0.1$ based on Least Significant Difference. ns – Non-significant at $p = 0.05$ . CV – Coefficient of variation.

When the diversity index data from ecosystems AQ, BL and PN were included in a regression analysis, arthropod diversity displayed a significant (*p<*0.05) positive linear dependence on plant diversity (Fig. 7). However, when the diversity data from PNEN were also included in the regression, a curvilinear dependence was observed (Fig. 8), where arthropod diversity decreased when vegetation diversity increased beyond a maximum.

**Figure. 7:**
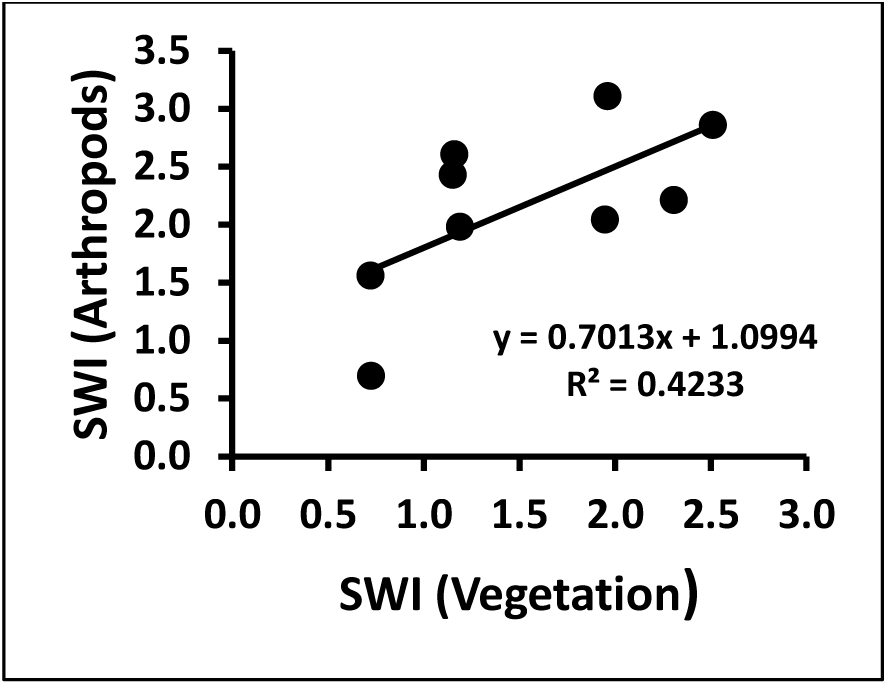
Dependence of arthropod diversity on vegetation diversity for Aquatic-based (AQ), Broad Leaved (BL) vegetations and *Pinus* monoculture (PN). SWI – Shannon-Wiener Index.

**Figure. 8:**
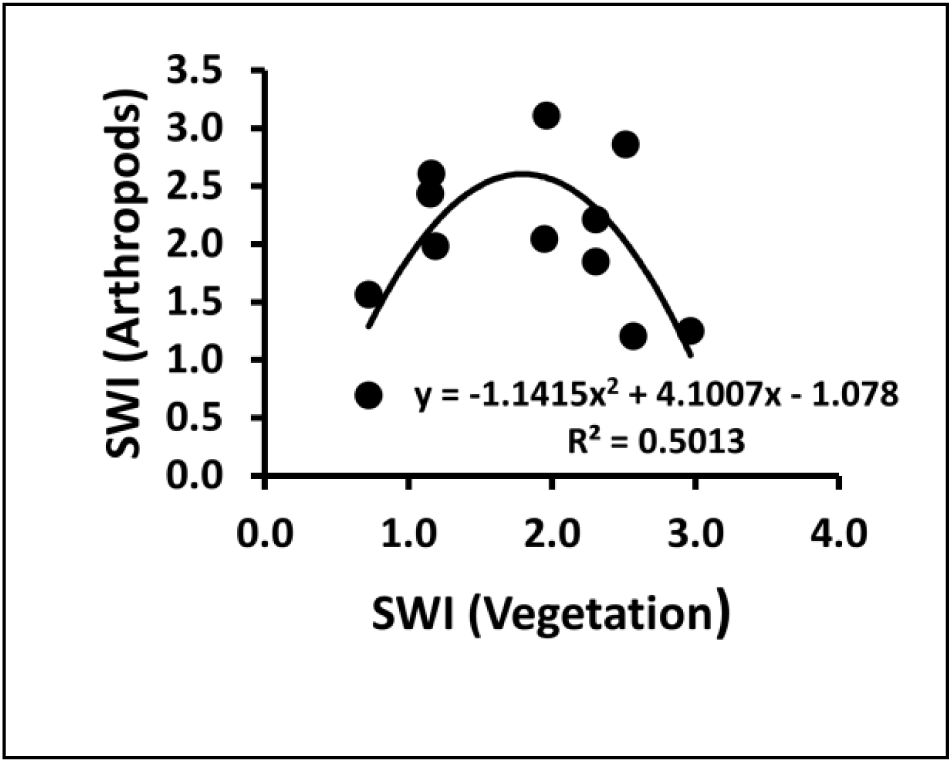
Dependence of arthropod diversity on vegetation diversity for all four ecosystems including enriched *Pinus* vegetation (PNEN). SWI – Shannon-Wiener Index.

According to the Eigen values of the Principle Component Analysis carried out to determine the variation among ecosystems based on the diversity of arthropod families, ecosystem AQ differed prominently from other ecosystems. This was because of the higher abundance of arthropods of orthopteran families such as Tettigonidae and Cicadellidae, coleopteran families such as Lampyridae, hemipteran families including Gerridae and Diapsididae and the lower abundance of several arachnid families including Pholcidae, Uloboridae, Scytodidae and Clubionidae (Fig. 9). Ecosystem BL, characterized by the high abundance of arthropods of Family Tettigonidae and low abundance of artrhopods of families Curculionidae also showed a clear separation from other ecosystems (Fig. 9). Based on their arthropod diversity, ecosystems PN and PNEN showed greater similarity to each other than to the other two ecosystems. This is primarily due to the high abundance of arthropod of families Curculionidae, Uloboridae and Pscheridae in PN and PNEN (Fig. 9). The PCA conducted based on vegetation diversity, the four ecosystems showed a highly prominent divergence (Fig. 10).

**Figure. 9:**
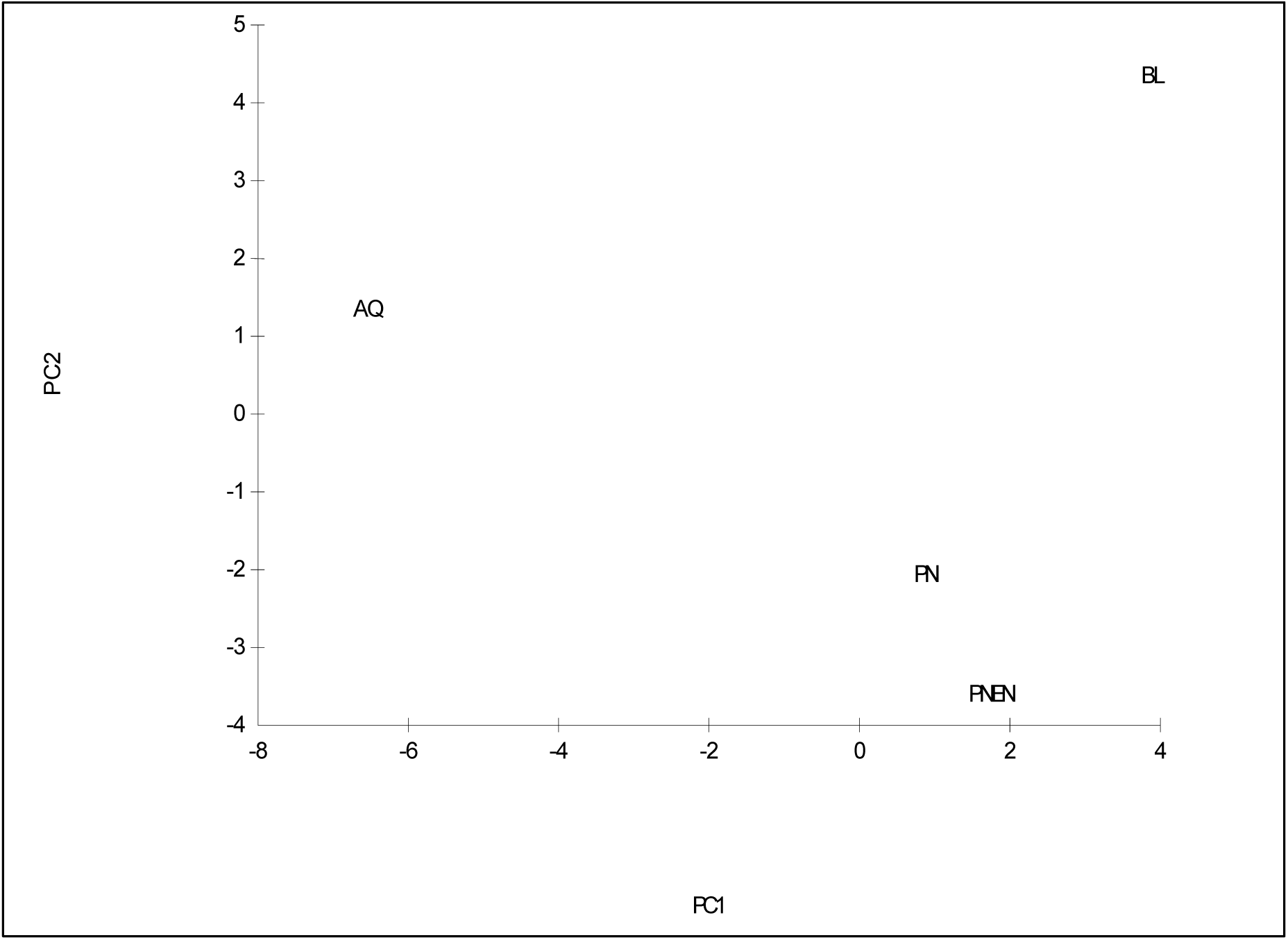
Variability of four ecosystems: Aquatic based vegetation (AQ), Broad leaved vegetation (BL), *Pinus* monoculture (PN) and enriched *Pinus* vegetation (PNEN) depending on the principal component analysis carried out on the diversity of arthropod families recorded

**Figure. 10:**
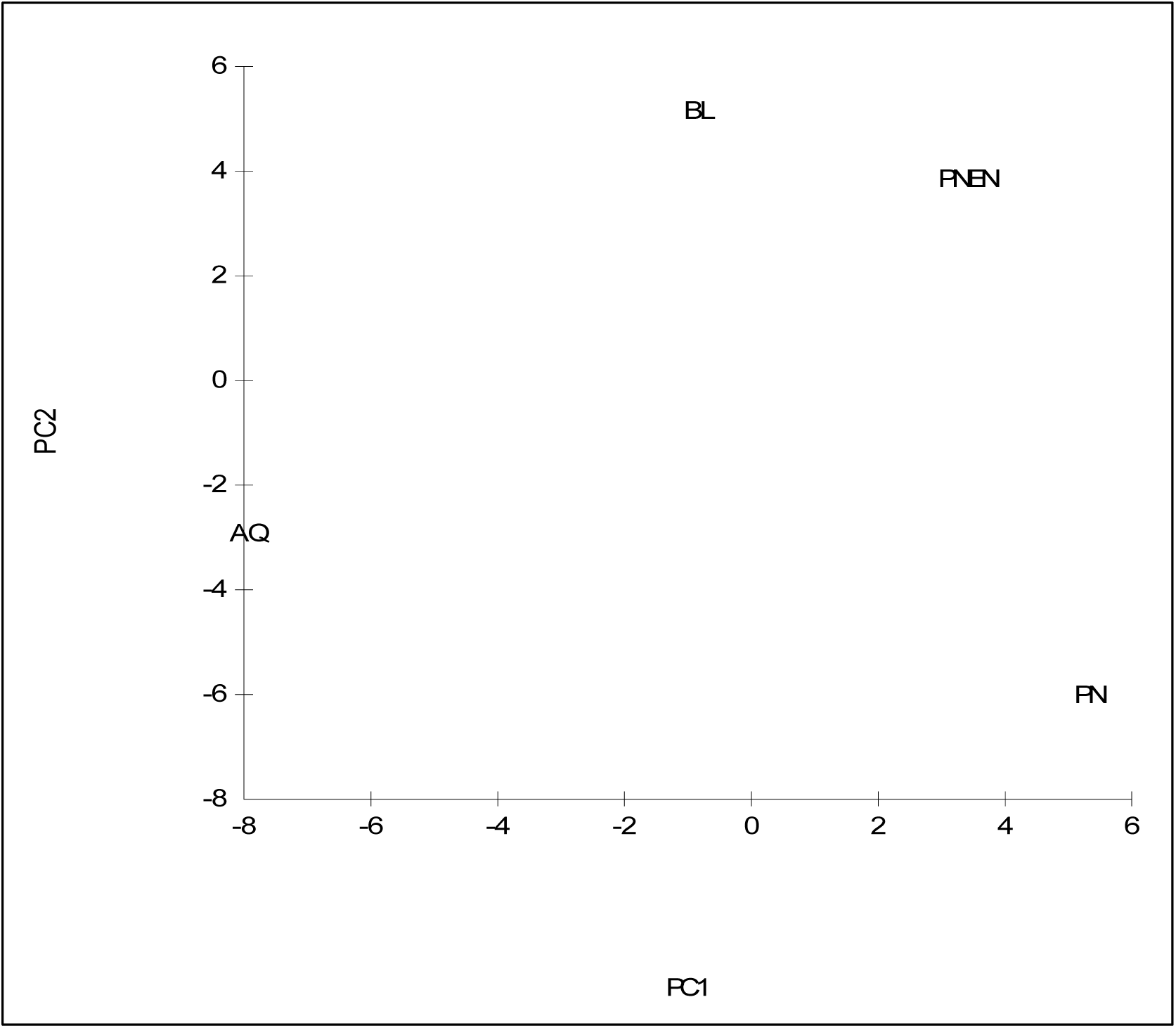
Variability among four ecosystems: Aquatic based vegetation (AQ), Broad leaved vegetation (BL), *Pinus* monoculture (PN) and enriched *Pinus* vegetation (PNEN) based on vegetation diversity

## Discussion

The principal focus of this work was to determine the dependence of arthropod diversity on vegetation diversity across four different ecosystems which are located close to each other in a mid-elevation humid tropical environment. Despite being a short-term study covering a limited area, our results provide important preliminary answers to the research questions that we posed at the beginning. Our results demonstrate significant variations in arthropod diversity (in terms of the Shannon-Wiener index) and their species richness among the four ecosystems, with the two *Pinus* based ecosystems (PN and PNEN) having significantly lower diversity than the two relatively un-disturbed ecosystems, the broadleaved evergreen forest (BL) and the aquatic-based environment (AQ). However, our results only partially confirmed the expected positive relationship between arthropod diversity and vegetation diversity, which had been demonstrated in previous work (Siemann *et al.*, 1998; Bangert *et al.*, 2005; Haddad *et al.*, 2009; Castangeyrol & Jactel, 2012). In agreement with such work, our results also showed that arthropod diversity across the three ecosystems which had not experienced direct and recent human intervention showed positive linear dependence on the vegetation diversity across the three habitats.

However, the most important finding of our work is the absence of an increase in arthropod diversity with the increased vegetation diversity in the enriched *Pinus* ecosystem even after two decades from artificial enrichment planting. This is in disagreement with the past work which had demonstrated the positive relationship between arthropod- and vegetation diversity. In fact such studies had involved artificial manipulation of the vegetation diversity and productivity via sowing seeds of different numbers of plant species (Haddad *et al.*, 2009), application of fertilizer at different rates (Siemann, 1998) and genetic hybridization (Bangert *et al.*, 2005). Contrary to the observed increases in arthropod diversity in the above work, enrichment planting and consequent increases in vegetation diversity, and most likely the net primary productivity (not measured in our work), in the enriched *Pinus* ecosystem had not increased the arthropod diversity. This finding indicates that the Resource Specialization Hypothesis (RSH) has a greater influence than the More Individual Hypothesis (MIH) in determining the arthropod diversity in the humid, tropical climate of the Hanthana mountain range. As the RSH is based on a majority of the arthropod species in a community showing host-specificity, it is likely that arthropod species specific to the introduced indigenous tree species have not been able to colonize and establish in the enriched *Pinus* ecosystem. This plausible when we take in to consideration the fact that many of the introduced tree species are not natives of the Hanthana mountain range. It is also possible that even after two decades, the environmental conditions within the enriched *Pinus* ecosystem are not conducive to support a broad diversity of arthropods species. Our observation that the dominant arthropod species in PNEN was a species of Family Formicidae supports this explanation because arthropods of Family Formicidae (ants) are a group which can adapt to any environmental condition rapidly.

Our observations that the dominant arthropod species in the aquatic and broadleaved evergreen forest environments are spider species confirms the fact that those ecosystems are subjected to minimum human intervention, because spiders are extremely sensitive to unfavorable environmental conditions (e.g. pesticides, inorganic fertilizer). This observation agrees with results obtained from parallel studies where highest species richness, species abundance, individual abundance and Shannon-Wiener index of spiders were recorded from natural forests of the Hanthana mountain range (Chathuranga & Ranawana, 2017). The dominance of spider species in two ecosystems having the highest plant diversity supports the conclusion of Haddad *et al.* (2009) greater vegetation diversity is conducive to development of a predator-dominated arthropod community.

The high-sensitivity of spiders to unfavourable environmental factors could be a reason for the spider population to be very low in the enriched *Pinus* ecosystem as it is an ecosystem exposed to inorganic fertilizer when the land was artificially enriched. The lower abundance of other arthropod species in ecosystems which are dominated by spider species can be due to the fact that spiders are voracious predators of insects. Nevertheless, this observation supports the fact that increased arthropod herbivores with increasing plant diversity increases diversity of arthropods at higher trophic levels, thus leading to a greater diversity of predators (Hunter & Price, 1992; Siemann *et al.*, 1998).

The dominant arthropod species in the *Pinus* ecosystem is *Dicaldispa* spp. commonly known as rice hispa. It is a pest infesting plants of the Family Poeceae (Grasses). This is plausible because the plant which shows the highest abundance in this environment is *Panicum maximum* (Family Poaceae), which is present as an undergrowth of the *Pinus* monoculture. Rice hispa could have been introduced to this ecosystem from the rice fields which are located in the neighboring areas.

We acknowledge the limitations of this study in that our findings are based on a single round of measurements on a limited number of small plots. Therefore, our findings can only be considered as preliminary and needs validation through a longer-term study involving a greater number of observational plots. Further research based on the foraging patterns and host-species relationships of the arthropods identified from these ecosystems is another promising research area which could explain the plant-arthropod interrelationships and community structure better. Nevertheless, we also point out that all four ecosystems that we used for sampling in our study have been free from any direct manipulations such as fertilizer application, selective thinning or enrichment. It has been so for the enriched *Pinus* ecosystem during the last 1 ½ decades. By using four different sampling methods, we have sampled the arthropods across a reasonable range of heights and depths. Therefore, our results constitute an important addition to the very limited knowledge-base on the dependence of arthropod diversity on vegetation diversity, especially in the humid tropical environments which the Hanthana mountain range represents. The regression models introduced from our work could also lay the foundation for more extensive studies aimed at describing community structures, interspecific relationships and finding pathways of conserving and enriching the biodiversity of this specific region, which is under severe pressure from human interference and urbanization.

